# Sexual differentiation of neural mechanisms of stress sensitivity during puberty

**DOI:** 10.1101/2020.12.02.408526

**Authors:** Emily C. Wright, Hannah C. Zakharenkov, Alexandra Serna Godoy, Alyssa A. Lake, Zhana D. Prince, Shwetha Sekar, Hannah I. Culkin, Pei X. Luo, Alison V. Ramirez, Tjien Dwyer, Amita Kapoor, Cody Corbett, Lin Tian, Andrew S. Fox, Brian C. Trainor

## Abstract

Anxiety disorders are more common in women than men, and this difference arises during puberty. Increased secretion of gonadal hormones during puberty influences brain structure and function, but the extent to which hormones modulate anxiety-related brain circuits is unclear. The slow developing California mouse (*Peromyscus californicus*) is an ideal species for studying the effects of hormones on brain function during adolescence. In adults social defeat stress reduces social approach and increases vigilance in females but not males. Here we show this sex difference is absent in juvenile mice, and that prepubertal castration sensitizes adult males to social defeat. Since adult castration has no effect on stress sensitivity, our data show that gonadal hormones act during puberty to program behavioral responses to stress later in life. In adults, calcium imaging in the medioventral bed nucleus of the stria terminalis shows that threatening social contexts increase calcium transients. Furthermore, prepubertal castration generalizes these responses to less threatening social contexts. Prepubertal treatment with the non-aromatizable androgen dihydrotestosterone acts in males and females to reduce sensitivity to social defeat in adults. Together, these data indicate activation of androgen receptors during puberty are critical for programing behavioral responses to stress in adulthood, highlighting a possible mechanism contributing to sex differences in anxiety.

**Significance Statement:** Puberty is a key period when sex differences in anxiety emerges. Gonadal hormone release increases during this time but it is largely unknown how they impact brain circuits and behavior. We show that androgens play a key role in programming behavioral responses to social defeat stress. The bed nucleus of the stria terminalis responds to social threats and these responses are more generalized in males without gonadal hormone exposure during puberty. Our findings highlight the importance of pubertal androgens in determining adult behavioral responses to social stress.

## Introduction

Adolescence is an important developmental period of cortical and sub-cortical re-organization (1–3) and a period when risk for anxiety disorders increases in girls (4–7). This increased risk is sustained into adulthood (8–13). The increased release of gonadal hormones may modulate these changes (14), as these hormones have important effects on brain structure (15–18) and function (19) by coordinating gene expression (20). Although human imaging studies demonstrate that neural circuits associated with anxiety are altered during pubertal development (21,22), little is known of how gonadal hormones alter stress-sensitivity of these circuits. This is important because stress exposure is an important risk factor for mental illness, and there is strong evidence in preclinical models that androgens can modulate behaviors related to anxiety or depression (23).

Social stressors exert powerful and sex-specific effects on anxiety-related behaviors in rodents (24,25). Female but not male adolescent rats exposed to a combination of social defeat and restraint stress showed reduced sucrose preference and increased passive coping in a forced swim test as adults (26). Adolescent social instability stress, in which rats are repeatedly housed with different cagemates, has stronger effects on anxiety-related behaviors in adult females than males (27,28). In adolescent mice, social isolation enhanced habit-related behaviors to a greater extent in females than males (29). A major outstanding question is whether pubertal hormones drive sex-specific behavioral and neurobiological responses to stress.

An important challenge for studying adolescent development in rodents is the relatively short temporal window. The most common domesticated mouse (30,31) and rat (32) lines are weaned at three weeks of age and are mature by five weeks of age. Here we introduce the California mouse (*Peromyscus californicus*) as a slow developing species that is ideal for studying adolescent development. The California mouse is an important model species for studying sex-specific effects of social stressors on anxiety-related behaviors (33). In adults social defeat increases the responsiveness of the bed nucleus of the stria terminalis (BNST) in adult females (34,35), which drives social avoidance and vigilance in novel social contexts (36–38). In adults these sex differences are independent of gonadal hormones (39,40). Here we use a combination of pubertal hormone manipulations, calcium imaging, and immunohistochemistry to demonstrate that androgens act during puberty to reduce behavioral responses to social defeat stress. These identify a neuroendocrine mechanism acting during puberty that directs developmental programing of stress-sensitive neural circuits.

## Results

### Late Onset of Pubertal Development in California mice

Complementary developmental measures suggest a slower developmental trajectory in California mice (Fig. 1A) compared to domesticated mice and rats. In males, testosterone levels increased with age (Fig. 1B, F_5,28_=6.51, p<0.01), but were not significantly elevated until post-natal day 90 (PN90, p<0.001, Cohen’s d=1.9). In females, progesterone levels increased with age (Fig. 1B, F_5.26_=2.56, p=0.05) but were not significantly elevated until PN70 (p=0.02, d=1.7). Other developmental measures such as gonadal weight, external genitalia, body weight, and pelage molt followed a similar trajectory (Fig. S1). Together, these data indicate that California mice have an extended period of development, ideal for studying adolescence.

**Figure 1:**
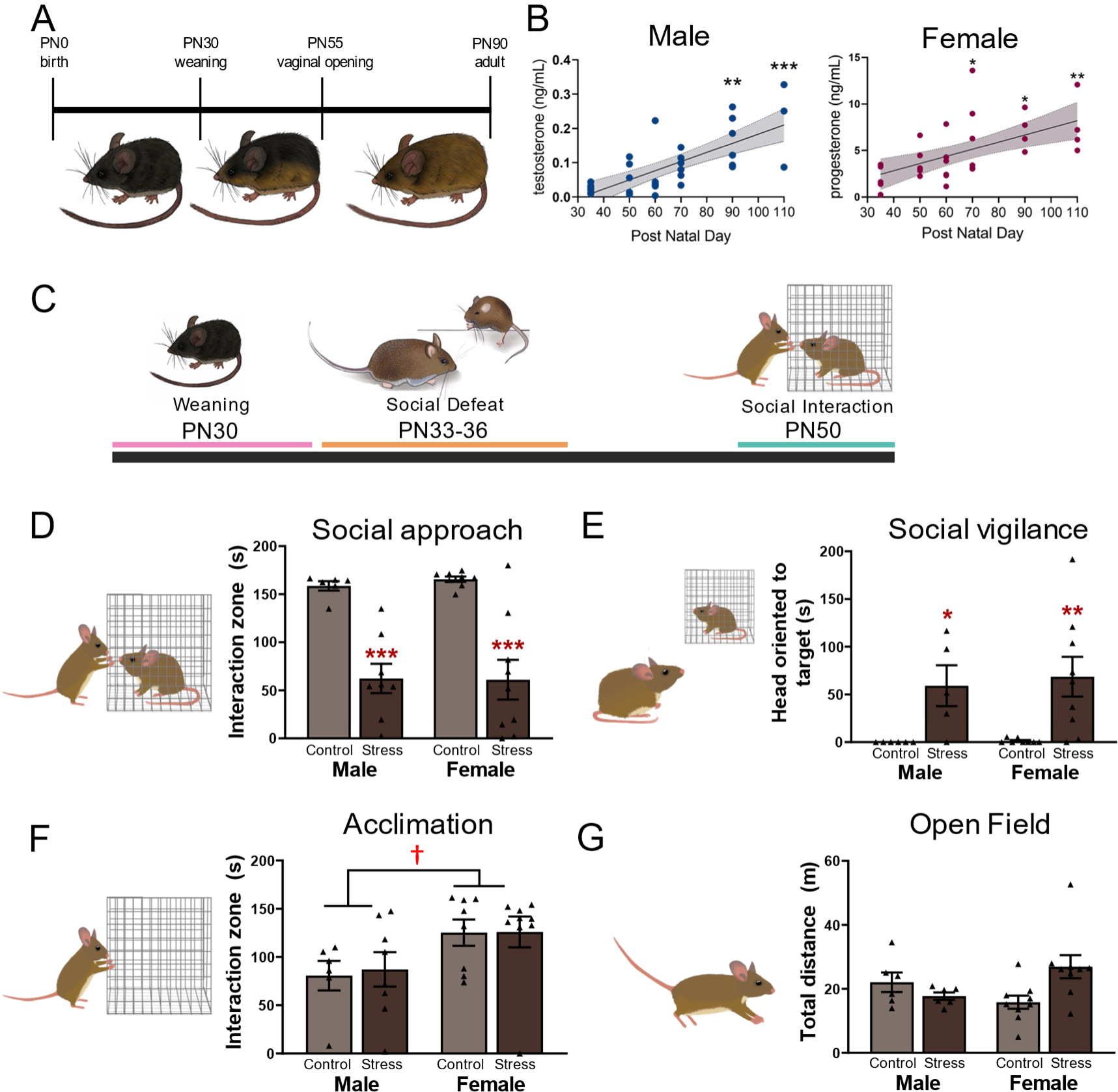
Delayed puberty in California mice is a period of sexual differentiation of behavioral stress responses. **A**. Summary of pubertal development in California mice **B.** In males testosterone levels are not increased until postnatal day (PN) 90 while in females progesterone levels increase at PN70 (n=4-5 per time point). **C.** Prepubertal California mice were exposed to social defeat or control conditions after weaning and tested as juveniles at PN50. **D,E.** Social defeat reduced social approach and increased social vigilance in both males and females (n=6-8 per group). **F.** Females investigated an empty cage during the acclimation phase more than males. **G.** There were no differences in locomotor behavior in the open field phase. *, **, *** p<0.05, p<0.01, p<0.001 vs. PN 35. *, **, *** p<0.05, p<0.01, p<0.001 vs same sex control. †p < 0.05 vs. males.

### Juvenile Male and Female California Mice Reduce Social and Approach and Increase Vigilance after Defeat

Juvenile mice were exposed to social defeat stress or control conditions after weaning and tested in a social interaction test as PN50 adolescents (Fig. 1C). Social defeat reduced social approach (Fig. 1D, F_1,27_=47.09, p<0.001) in both males (p<0.001, Cohen’s d=3.0) and females (p<0.001, d=2.4). Defeated also increased social vigilance (Fig. 1E, Kruskal-Wallis=16.33, p<0.001) in both males (p=0.03, d=1.6) and females (p=0.02, d=1.8). These results contrast with observed sex-differences in adults, which show that social defeat decreases social approach (35,39–41) and increases vigilance (42) in females but not males. During the acclimation phase females spent more time investigating an empty cage than males (Fig. 1F, F-_1,27_=6.64, p=0.016) with no effects of stress. There were no differences in locomotor behavior in the open field phase (Fig. 1G), time spent in the center (Fig. S2A) or vigilance behavior during the acclimation phase (Fig S2B). Together with previous work, these results suggest that developmental changes during puberty contribute to sex differences in stress responses in adults.

### Gonadal Hormones Reduce Susceptibility to Defeat in Adult Males

To assess the impact of gonadal hormones during puberty, males were assigned to prepubertal castration, sham surgery, or no surgery (Fig. 2A). Gonadectomy affected social approach only in males that were exposed to social defeat as adults (Fig. 2B, trt*stress F_2,41_=3.32, p<0.05). Castrated males exposed to defeat had reduced social approach versus controls (p<0.0001, d=2.6) whereas there was no effect of defeat in sham or no surgery males. Similarly, social defeat increased vigilance in castrated (Fig. 2C, Mann-Whitney U=23.5, p<0.01, d=1.6) but not sham or no surgery mice. There were no differences in behavior during the acclimation (Fig. 2D, S3A) or open field phases (Fig. 2E, S3B). The results contrast sharply with previous results that showed no effect of adult castration or ovariectomy on stress-induced social avoidance in California mice (40), and point to gonadal hormones acting during adolescence as a key mechanism of sexual differentiation. These data suggest that pubertal hormones may play a protective role in organizing the brain to diminish the effects of social defeat on social approach and vigilance. The BNST plays a key role in modulating stress-induced social avoidance and vigilance in males and females (35,38,42). Since prepubertal castration did not affect behavior in unstressed mice, we used fiber photometry to assess effects of castration on BNST neural activity in males exposed to defeat.

**Figure 2:**
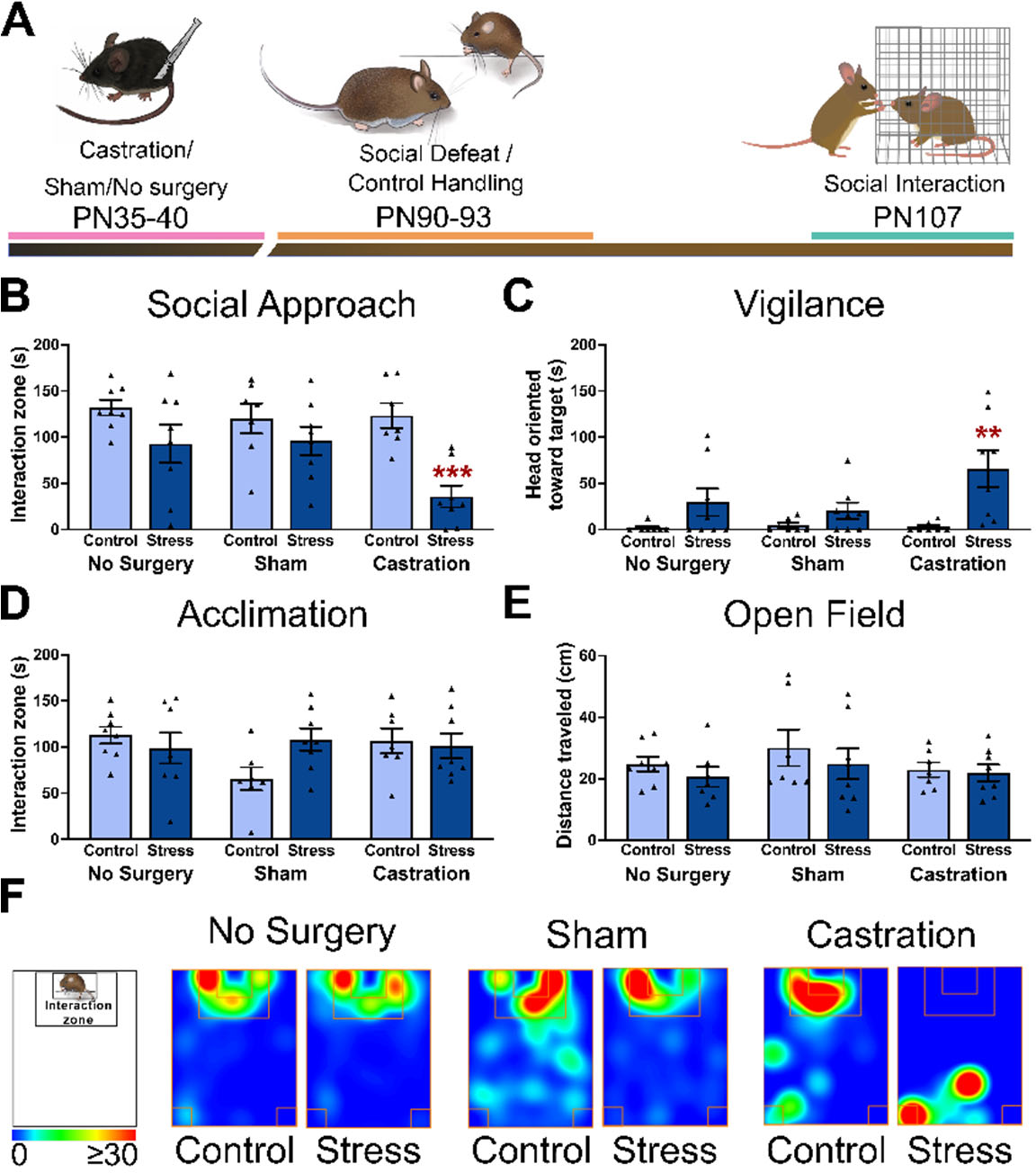
Prepubertal castration increases sensitivity to social defeat in adulthood. **A.** Timeline of prepubertal castration and behavioral testing in adults. **B,C.** Social defeat decreased social approach and increased social vigilance in castrated males but not sham or no surgery controls (n=7-8 per group). **D,E.** No differences were observed during the acclimation of open field phase. **F.** Representative heatmaps for the interaction phase showing reduced time spent in the interaction zone castrated males exposed to social defeat. **, *** p<0.01, p<0.001 vs control.

### Calcium imaging of the BNST activity in castrated and intact males

The BNST modulates social approach and avoidance (34,42), so we performed calcium imaging of GCaMP6f in medioventral BNST to determine how prepubertal castration impacted neural responses in novel social contexts (Fig. 3A, 3B). Previous immediate early gene analyses show that social stress increases activity within the BNST (43,44), but does not have the temporal resolution to determine whether increased activity occurs during brief episodes of social approach or more extended periods of avoidance. Our results indicated that GCaMP signals coincided with social contact with target mice rather than avoidance, regardless of gonadal status. Using DeepLabCut (Fig. 3C, 3D) we determined orientation (45) and distance between focal mice and non-aggressive or aggressive target mice. Increased ΔF/F in BNST was observed when focal mice were within one body length (8 cm) of target mice and within the central (Fig. 3D, 0°≥40°, β= - 0.438, z= −9.783, p>0.001) or peripheral visual field (40°≥100°, β= −0.486, z= −10.356, p>0.001), but not if the focal mouse was facing away from target mice (100°≥180°). We then examined ΔF/F in specific behavioral contexts.

**Figure 3:**
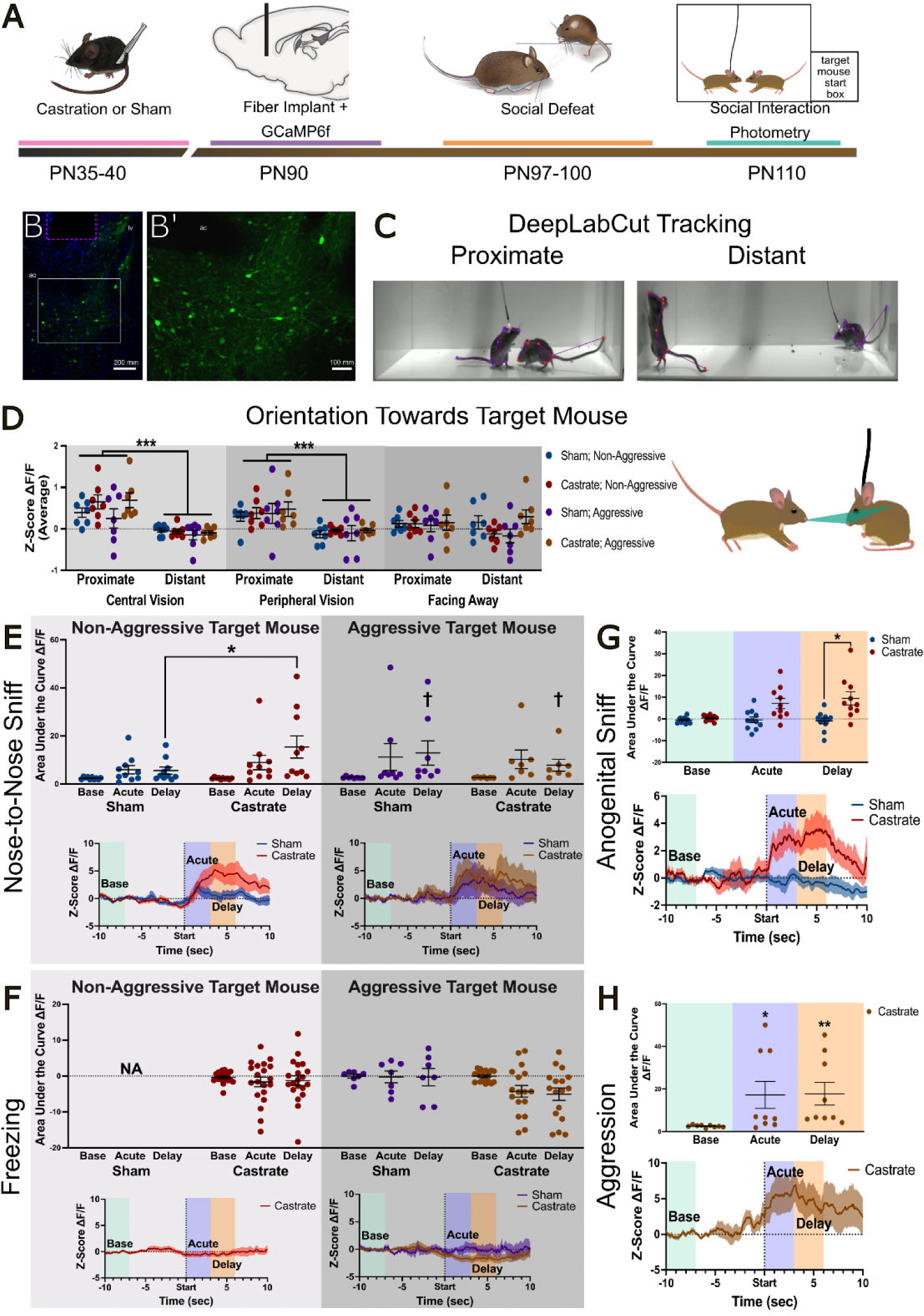
Prepubertal castration increases BNST neural activity in sensitivity to social defeat in adulthood. **A.** Experimental timeline for fiber photometry observations of GCaMP6f in the ventral BNST of prepubertally castrated or sham surgery California mice. **B.** Photomicrographs of GCaMP in BNST at low (B) and high (B’) magnification. The magenta box indicates position of the fiber. **C.** Representative images showing DeepLabCut tracking of mice in proximate and distant conditions. **D.** DeepLabCut tracking of body midpoint and nose of the focal mouse was used to determine the orientation of the focal mouse to the target mouse nose (green triangle). GCaMP6f signals were significantly stronger when the focal mouse was within 8 cm of the target mouse but only when the target mouse was in central vision (0°≥40°) or peripheral vision (40°≥100°). **E.** Nose-to-nose sniffing with a non-aggressive target mouse induced a delayed increase in ΔF/F in castrated but not sham males. In contract nose-to-nose sniffing with an aggressive target mouse increased ΔF/F in both sham and castrated males. **F.** There were no changes in ΔF/F following bouts of freezing. **G.** Anogenital sniffing of non-aggressive target mice increased ΔF/F in castrated but not sham males. **H.** When castrated males were attacked by aggressive target mice, increased ΔF/F was observed. Sham males were not attacked enough for analysis. * p< 0.05, **p<0.01 vs sham. *** p<0.001 vs proximate, †p<0.05 vs. baseline. ac= anterior commissure.

Intriguingly, the most robust changes in ΔF/F occurred after close contact with an aggressive target mouse. After engaging in nose-to-nose sniffing, both castrated and sham males showed a delayed increase in BNST ΔF/F (Fig. 3E, β=10.22, z=2.05, p=0.04). In contrast, when engaging in nose-to-nose sniffing with a non-aggressive target mouse, castrated males showed a larger increase in BNST ΔF/F than sham males (Fig. 3E, β=13.2, z=2.23, p=0.03). These results suggest that BNST ΔF/F is enhanced by social threats and that this response is more generalized in prepubertally castrated males. We also examined BNST following bouts of freezing, which was robustly induced by aggressive targets (Fig. S4) while non-aggressive targets induced more freezing in castrated males than shams (t_11_=2.43, p=0.03). There were no acute or delayed changes in ΔF/F following bouts of freezing (Fig. 3F). We next examined bouts of anogenital sniffing, which were less frequent and limited to non-aggressive target mice (Fig. S4). Similar to nose-to-nose sniffing castrated mice showed a larger delayed increase in ΔF/F in BNST than shams (Fig. 3G, β=6.12, z=2.06, p=0.04. Finally, when castrated males were attacked by aggressive target mice, there were acute (Fig. 3H, β=5.71, z=2.56, p=0.01) and delayed (β=5.71, z=2.67, p=0.008) increases in ΔF/F. Together these data suggest that neuronal activity in BNST increases after potentially-threatening social situations, and is generalized to less threatening contexts after prepubertal castration. To assess the extent to which prepubertal castration impacted neuronal activity in other circuits modulating social approach, we used c-fos immunohistochemistry.

### Gonadal Hormones Reduce Neural Activity in PVN Oxytocin Neurons

Social defeat stress has been demonstrated to elevated the activity of oxytocin releasing neurons in PVN (34), a response that has been associated with heightened activation of stress-reactive neurons expressing oxytocin receptors in the anteromedial BNST (42). We used oxytocin/c-fos immunohistochemistry (Fig. 4A,B) to examine anterior and posterior PVN, which differ in connectivity (46) and stress sensitivity (34). In anterior PVN, prepubertal castration increased oxytocin/c-fos colocalizations when compared to sham (Fig. 4C’, Mann-Whitney U=38, p=0.01) in mice that had undergone social defeat. Oxytocin/c-fos colocalizations were negatively correlated with social approach (Fig. 4C”, Spearman ρ=-0.55, p<0.01) and positively correlated with social vigilance (Fig. S5A, ρ=0.59, p<0.01). In the posterior PVN castration increased oxytocin/c-fos colocalizations regardless of stress status (Fig. 4D’, Mann-Whitney U=113.5, p=0.04) and oxytocin/c-fos colocalizations was negatively correlated with social approach (Fig. 4D”, ρ=-0.41, p=0.04) but not vigilance (Fig. S5B). We also examined the effects of prepubertal castration on c-fos in the anteromedial BNST, where oxytocin induces social avoidance and vigilance (38). Prepubertal castration increased the number of c-fos positive neurons in BNSTam (Fig. 4E’, S5D, F_1,21_=4.51, p=0.046), regardless of stress status. The number of c-fos cells was negatively correlated with social approach (Fig. 4E”, ρ=-0.46, p=0.02) but not vigilance (Fig. S5C).

**Figure 4:**
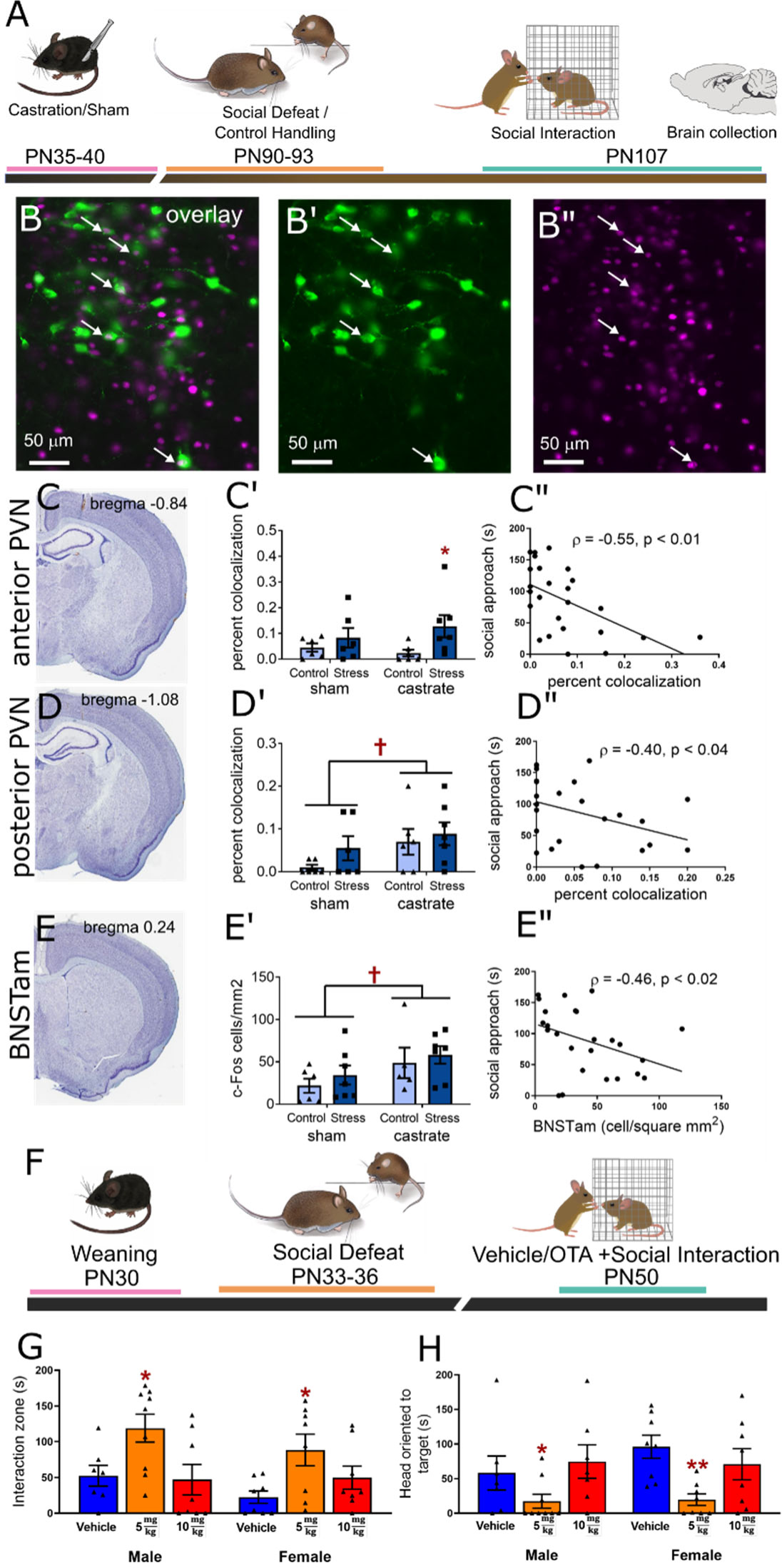
Sexual differentiation of oxytocin-dependent circuits of social avoidance occurs during puberty. **A.** Experimental timeline for immunohistochemistry analyses in prepubertally castrated or intact California mice. **B.** Overlay of oxytocin (green) and c-fos (magenta) immunostaining in the paraventricular nucleus. Arrows indicate colocalizations in oxytocin (B’) and c-fos (B”) images. **C.** In the anterior PVN social defeat increased colocalizations (n=6-7 per group) in prepubertally castrated males but not intact males(C’) and that social approach was negatively correlated with oxytocin/c-fos colocalizations (C”). **D**. In the posterior PVN castration increased oxytocin/c-fos colocalizations regardless of stress status (D’) and social approach was negatively correlated with colocalizations. E. In anteromedial BNST, where oxytocin receptors drive social avoidance, prepubertal castration increased c-fos immunoreactivity regardless of stress status (E’) and c-fos positive cells were negatively correlated with social approach (E”). **F**. Experimental timeline for examining effects of oxytocin receptors on social behavior. **G,H**. In both males and females exposed to social defeat an i.p. injection of 5 mg/kg of the oxytocin receptor antagonist L368,899 increased social approach and decreased social vigilance (n=7-9 per group). *, ** p<0.05, control/vehicle. †p < 0.05 vs. intact.

We also tested whether defeat induced social avoidance and vigilance in adolescent mice was dependent on oxytocin receptors (Fig. 4F) as in adults. In both males and females, treatment with 5 mg/kg i.p. of the oxytocin receptor antagonist L-368,899 30 min before testing increased social approach (Fig. 4G, main effect of dose F_2,42_=8.42, p<0.001) and decreased vigilance (Fig. 4H, Kruskal-Wallis=15.36, p<0.009). There were no differences in behavior during the acclimation (Fig. S5E,S5F) or open field (Fig. S5G, S5H) phases. Together these results suggest that stress-induced social avoidance and vigilance in both male and female juvenile mice is dependent on oxytocin receptor activation, as seen in adult females. Additionally, the results indicate that male pubertal hormones program these circuits to be less active in adulthood.

### Androgens Reverse the Effects of Prepubertal Castration on Behavior

To test the impact of androgen replacement at puberty, juvenile males were prepubertally castrated and randomly assigned to receive a silastic implant containing testosterone, the non-aromatizable androgen dihydrotestosterone (DHT), or sealant only (Fig 5A). These implants produce plasma hormone levels within physiological range of adult male California mice (47). At PN90, mice were exposed to defeat stress and then tested in a social interaction test. Results demonstrated that hormone replacement altered social approach (Fig. 5B, F_2,22_=6.2, p<0.01) and vigilance (Fig. 5C, Kruskal-Wallis=13.13, p<0.001). For social approach both testosterone (p=0.002, d=1.7) and DHT (p=0.002, d=1.0) treatment yielded significantly higher levels of social approach compared to males treated with empty implants. Similarly for vigilance both testosterone (p<0.001, d=3.1) and DHT (P=0.049, d=1.1) treated males had lower vigilance than males treated with empty implants. There was a nonsignificant trend for testosterone treated males to have lower vigilance than DHT treated males (p=0.078, d=1.0). During the acclimation phase T and DHT treatment increased approach to an empty cage (Fig. S6A, F_2,22_=5.02, p<0.02) and there were no differences in behavior during the open field phase (Fig. S6B, S6C). We also tested whether DHT treatment at puberty could impact female behavior (Fig. 5D). In social interaction tests performed before stress exposure, there were no differences in social approach (Fig. 5E), social vigilance (Fig. 5F), acclimation (Fig. S6D) or open field behavior (Fig. S6E,F). Importantly, after social defeat, hormone treatment altered social approach (Fig. 5E, F_2,23_=3.76, p= 0.04) with DHT increasing social approach versus empty implants (p=0.02, d=0.1). Effects on vigilance were weaker (Kruskal-Wallis=5.04, p=0.08), with a post-hoc test indicating lower levels of vigilance in DHT treated mice versus empty implant (Mann-Whitney=11, p=0.028, d=1.2). After stress exposure DHT treatment increased approach to an empty cage (Fig. S6D, F_2,23_=5.32, p= 0.01) while there were no differences in the open field phase (Fig. S6E, S6F)

**Figure 5:**
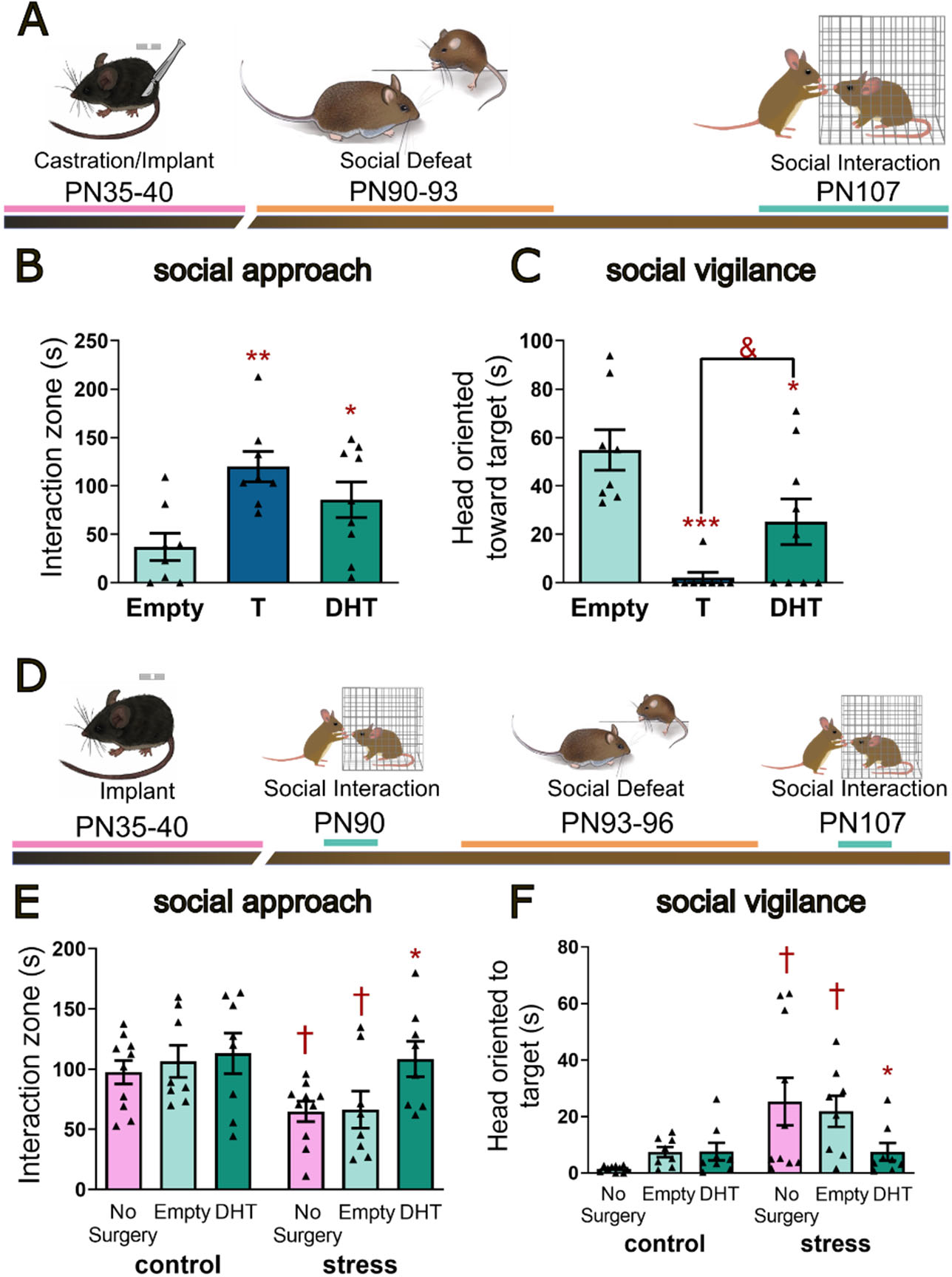
Androgen treatment at puberty reduces effects of social defeat on social approach and vigilance in males and females. **A.** Experimental timeline for surgery, social defeat, and social interaction testing in males. **B**. Castrated males treated with dihydrotestosterone (DHT) or testosterone (T) implants had higher social approach than males treated with empty implants (n=7-8 per group). **C**. Treatment with DHT or T also lowered social vigilance. **D**. Experimental timeline for surgery, behavior testing, and social defeat in female California mice. **E**. Hormone treatment had no effect on social approach before stress exposure but after stress exposure DHT treated females showed more social approach than females treated with empty implants or no surgery controls (n=8-10 per group). **F**. Similarly there were no differences in social vigilance in before stress but after stress social vigilance increased in no surgery control and empty implant females but not DHT females. *p<0.05, ** p<0.01, *** p<0.001 vs. empty implant. & p<0.05 vs T. †p < 0.05 vs. control (pre-stress).

## Discussion

Sex differences in stress-sensitivity emerge at puberty in both humans (48,49) and animals (26–28), coinciding with an increased incidence of anxiety in women (4,6,7,12). However, little is known about the underlying mechanisms. Using the California mouse, we found stress-induced social avoidance and vigilance occurs in both sexes before puberty. Prepubertal castration enhanced behavioral responses to stress in adult males. Gonadal hormones do not affect social avoidance in adults (40,50), suggesting that puberty is a critical period for sexual differentiation. Dihydrotestosterone replacement during puberty in castrated males and intact females reduced stress-induced social avoidance in adulthood, highlighting an essential role for androgen receptors. Using real-time calcium imaging we found that neurons in ventral BNST are highly responsive to threatening social contexts, and that prepubertal castration generalized these responses to non-aggressive mice. Our findings indicate that androgens play an organizational role during puberty, which attenuates behavioral and neural responses to social stress in adulthood.

### BNST neurons respond to social contact

Immediate early gene analyses in rodents (34,51–54) and neuroimaging studies in primates (55) and humans (56) show that stressful social contexts increase neural activity within the BNST. However, these analyses lack the temporal resolution to associate changes in neural activity with specific behaviors. Our calcium imaging data show that ventral BNST calcium transients occurred after close contact with target mice. Specifically, increased activity was observed following active engagement with aggressive target mice, such as nose-to-nose sniffing or aggression. Intriguingly no increases occurred during freezing, a behavior that does not involve active social engagement. While it has been hypothesized that the BNST encodes more distant threats (57), our results are more consistent with recent hypotheses that the BNST is highly responsive to proximate threats (58). The strength of calcium transients in the presence of non-aggressive target mice was dependent on exposure to pubertal hormones.

Pre-pubertally castrated males exhibited more freezing and heightened BNST calcium transients in the presence of non-aggressive target mice. These results are consistent with our findings showing that pre-pubertal castration leads to increased avoidance and vigilance towards unfamiliar target mice, similar to stressed adult females (40,59). These findings suggest that the absence of male gonadal hormones during puberty may alter valence assessment of novel social contexts (60,61), potentially leading to an over-generalization of threat, which is a key characteristic of anxiety disorders (62). Our results align with imaging studies in nonhuman primates (63) and humans (64) reporting stronger BNST responses in individuals with elevated anxiety-related behaviors. Intriguingly, androgens can reduce neuronal excitability in both adult (65) and adolescent rodents (66–68). These findings are in line with our observations that prepubertal castration led to increased c-fos/oxytocin colocalizations in the PVN and c-fos expression in anteromedial BNST. Taken together, our results suggest that androgens may play a role during puberty to reduce the excitability of circuits that are affected by social defeat.

The question remains as to which BNST cell types respond to social threats. This is further complicated by the fact that the BNST exhibits heterogeneity in neural responses, even within genetically defined cell populations (69–71). This variation could be explained by projection-specific populations within a cell-type (72). This is supported by optogenetic studies showing the projection-specific effects of BNST neurons on behavior (73–76). Future research considering genetic and projection-specific populations during neuronal recording of activity in different social contexts should be highly informative.

### Activation of androgen receptors during puberty blunts stress induced social avoidance

In adult hamsters, dominant males have more androgen receptor expression in the medial amygdala (MEA) than subordinates and pharmacological inhibition of AR in the MEA of dominant males increases sensitivity to social stress (77). Similarly testosterone can exert acute anxiolytic effects in a variety of behavioral assays (78–80). Interestingly, in adult California mice, androgens did not appear to affect social approach regardless of stress exposure (40). Here we show that androgens act during puberty to influence how social behavior is impacted by stress, suggesting that even in species where androgens do not have an overt role in regulating stress-sensitivity in adults, androgens can program behavioral responses during adolescence. This may be particularly relevant for humans, where the relationship between gonadal hormones and stress responses in adults can be inconsistent (81–83). Although adolescent DHT and testosterone treatment had similar effects on social approach in stressed males, the effect of DHT on vigilance was weaker. Similar effects were observed in females. These results suggest that estrogen receptors may play a complementary role in pubertal organization of stress-induced behavior. Estrogens can enhance anxiety-related behaviors by activating *Esr1* or exert anxiolytic effects via *Esr2* (84). Both receptors are present in the BNST, and future studies should consider the possible developmental effects of these receptors on stress-induced social vigilance.

## Functional implications

Steroid hormones alter brain development during puberty (15,17,19), yet how these changes affect anxiety-related behavior has been unclear. We showed that sex differences in social stress effects on behavior are driven, in part, by pubertal androgen exposure in males. A lack of pubertal hormone exposure increased reactivity of BNST neurons during social engagement. Interestingly, the effects pubertal androgens on social behavior were only apparent following exposure to social defeat. This demonstrates that organizational effects of pubertal hormones can cause latent vulnerabilities that are only revealed after stressful social experiences. Our results suggest that it will be worthwhile to consider whether testosterone levels during adolescence predict behavioral or neural responses to stressors in adulthood. Overall, our research sheds light on how androgens shape the complex interplay between brain circuits and behavioral sensitivity to stress.

## Methods

### Animals

All studies were conducted with California mice (*Peromyscus californicus*) raised in a colony at UC Davis. Mice were housed in same sex groups (2-4) in clear polypropylene cages with Sani-Chip bedding (Harlan Laboratories, Indianapolis, IN, USA), Nestlets (Ancare, Bellmore, NY, USA) and Enviro-Dri (Eco-bedding, Fibercore, Cleveland, OH, USA). Mice were kept on a 16L:8D light cycle and had *ad libitum* access to food and water. All procedures were approved by the Institutional Animal Care and Use Committee at the University of California, Davis and in accordance with National Institutes of Health (NIH) guidelines.

### Puberty Quantification

For measures of first vaginal opening, preputial separation, weight, and coat color mice were briefly anesthetized (>1 minute isoflurane) before being weighed, photographed, and assessed for vaginal opening/preputial separation. A separate set of mice were euthanized to determine uterine or testes size. Trunk blood was collected and plasma obtained and frozen for hormone analyses. Plasma was extracted and hormones quantified on a QTRAP 550 quadruple linear ion trap mass spectrometer equipped with an atmospheric pressure chemical ionization source (LC-MS/MS) as previously described (85). Quantitative results were recorded as multiple reaction monitoring (MRM) area counts after determination for the response factor for each hormone and internal standard. We also quantified the transition from the juvenile pelage (dark gray) to adult (brown) adult pelage (86) by quantifying digital images in ImageJ as previously described (87).

### Juvenile Social Defeat

Male and female juveniles (PN34-36) underwent 3 consecutive days of social defeat stress (or control handling) as described previously (50). During each episode of defeat the focal mouse was placed in the home cage of a novel male-female adult resident pair. The opposite sex resident mouse was removed from the cage prior to the start of the defeat session. Each session of defeat lasted for 7 minutes, or until the test mouse received 7 bites. Control mice were placed into a clean, empty cage for 7 minutes across 3 consecutive days.

On PN50 mice underwent a social interaction test. The social interaction test had 3 phases: open field, acclimation, and interaction. In the open field phase, the mouse was placed into an open area and allowed to explore it for 3 minutes. In the acclimation phase, an empty cage was placed at one end of the arena and the mouse was allowed to explore for 3 minutes. In the interaction phase, a caged, unknown adult conspecific replaced the empty cage at one end of the arena, and the mouse was allowed to explore for 3 minutes. Time in the interaction zone (8 cm from the cage) is defined as “social approach” was scored with AnyMaze software (40). Duration of vigilance behavior was hand scored from a video recording. Vigilance was defined as any time the test mouse was sitting still, head oriented toward the target mouse, while outside of the interaction zone (37).

### Prepubertal Castration

Male juveniles were randomly assigned to be castrated, sham surgery, or no surgery control between PN35 and PN40 (47). During castration mice were anesthetized with isoflurane and treated with 0.1mg/kg buprenorphine and 5mg/kg of carprofen. Non-surgery controls were included to control for possible effects of early life exposure to the anesthesia isoflurane (88). These mice received no manipulations until the start of social defeat stress/control handling. At PN90-92 all mice underwent 3 days of either social defeat or control handling. At PN106 mice were tested in a social interaction test.

### Calcium imaging of the BNST activity in castrated and intact males

Male juveniles were randomly assigned to castration or sham surgery after weaning. As adults, all mice received an injection of AAV9.Syn.GCaMP6s at a rate of 100nL/min for a total volume of 500nL in the ventral BNST (AP +0.45mm, ML +/-1.0mm, DV −5.6mm). The virus was allowed to diffuse for 10 minutes before the needle was withdrawn. An optical fiber (Doric) with 2.5 mm core and 0.66 NA threaded through a ceramic ferrule was implanted at the injection site and the ferrule was secured to the skull with a layer of C&B Metabond (Parkell), followed by a layer of dental cement to form a thick headcap. Mice were housed two per cage with a clear, perforated acrylic divider that allowed auditory, tactile and olfactory contact.

Mice recovered for 1 week before undergoing 3 consecutive days of social defeat stress. Ten days later mice began 3 consecutive days of patch cord habituation in which the patch cord was gently coupled to their optical fiber implant. The mouse then explored a novel cage for 10 min. The next day mice were tested in a social interaction test with freely moving target mice. The focal mouse was placed into an empty area (51×25.4×76 cm) attached to a small box (13×10×18 cm) with a sliding door for 6 min. Photometry recording was performed with an isosbestic channel of 405nm and an excitatory channel of 470nm, both set to 50μW output. Photometry data from the acclimation period were excluded from analysis due to initial bleaching. Next, an unfamiliar, nonaggressive adult male target mouse was introduced into the arena through the sliding door. Mice were allowed to freely interact for 3 min. The target mouse was removed and then a sexually experienced, aggressive male (from a previous social defeat episode) was introduced into the arena for 3 min. After testing brains were collected to confirm viral expression and placement of the fiber.

We used Deeplabcut to quantify distance and orientation of the focal mouse in relation to target mice with precision that matched the high temporal resolution of photometry data (89– 91). Video was taken from the side view at a rate of 30fps. And training was performed to track the nose, right ear, left ear, body midpoint, front leg, back leg, tail base, and tail tip for both the test mouse and stimulus mouse. The tracking for the nose and body midpoint were used in order to determine proximity, defined as times when the body midpoint of the two mice were within 8 cm (the approximate length of a California mouse) of each other. We determined the direction the test mouse was facing in relation to the stimulus mouse using their coordinate points obtained from the deeplabcut tracking (test mouse nose, body midpoint, and target mouse nose). We binned angles as within the test mouse’s central vision (0°≥40°), peripheral vision (40°≥100°), or facing away from the stimulus mouse (100°≥180°) (45). All scripts are deposited at https://github.com/bctrainorlab/behavioral_quantification. Nose to nose sniffing, anogenital sniffing, freezing and aggressive behavior directed towards focal mice were scored by an observer without knowledge of the treatment group using BORIS.

### Pubertal hormone manipulation studies

Male juveniles were castrated and received a subcutaneous silastic implant (i.d. 0.04 in, o.d. 0.085 in) containing 1mm crystalline testosterone, dihydrotestosterone, or Dowsil Sealant (3145 RTV MIL-A-46146). These implants yield hormone levels that are within the physiological range of intact male California mice (92). At adulthood mice underwent 3 days social defeat and were tested in a social interaction test two weeks later.

Intact females were randomly assigned to receive a DHT or empty implant at PN35-PN40 as described for males. At PN90 all mice were tested in a social interaction test. Three days later females were exposed to 3 days of social defeat on consecutive days. One week later females were tested again in a social interaction test. This within-subjects design produces behavioral effects similar to the between-subjects design used for males (93).

### Immunohistochemistry

Sections from castrated or intact males were stained for either oxytocin/c-fos (PVN) or c-fos (anteromedial BNST) as previously described(34). Sections were blocked in 10% normal goat serum for 20 minutes and then incubated in rabbit anti-cfos (Synaptic Systems, 1:K) in PBS 0.5% TritonX overnight at room temperature. For oxytocin double staining, sections were washed in PBS and then incubated in goat anti-rabbit AlexaFluor 555 (Molecular Probes, 1:500 in PBX-TX) for 2 hrs. Sections were then incubated in mouse anti-oxytocin (Sigma-Aldrich MAB5296 1:2000 in PBS-TX) overnight at 4 C. Sections were then washed in PBS and incubated goat anti-mouse AlexaFluor 488 (Molecular Probes, 1:500 in PBS-TX) for 2 hrs. For BNSTam sections, sections were washed in PBS and incubated in biotinylated goat anti-rabbit (Vector labs, 1:500 in PBS-TX for 2 hrs). Sections were washed in PBS and then incubated in avidin-conjugated peroxidase (Vector labs) for 30 min. After washing in PBS sections were developed in nickel enhanced diaminobenzidine (Vector labs) for 2 min.

### Juvenile Social Defeat + OTA Injections

Male and female juveniles (PN34-36) underwent 3 consecutive days of social defeat stress. At P50 mice were tested in a social interaction test. 30 minutes before the start of the social interaction test, each mouse received an intraperitoneal (i.p.) injection of 5mg/kg or 10mg/kg of the oxytocin antagonist (OTA) L-368,899 (37), or an injection of sterile phosphate buffered saline (PBS).

### Statistics

Statistical analyses for experiments (excluding fiber photometry experiments) were performed using R statistical software. Normality was assessed via QQPlot. A Flinger-Killen test was used to assess homogeneity of variance. Steroid hormone data were log transformed for analysis. Hormone and most behavioral and cell count data were analyzed using ANOVA with planned comparisons for comparing treatment groups with controls. Social vigilance and oxytocin/c-fos colocalization data had heterogeneous variability, so non-parametric analyses were used.

Photometry data was analyzed using a custom Python script, down sampled to 30 samples per second to match the 30fps video framerate. 405nm was used as the isosbestic wavelength and 470nm as the excitatory wavelength. To correct for motion artifact and bleaching of the fluorophore, the 405nm signal was fit to a biexponential model and subtracted from the excitatory output. Such that ΔF/F = [100*((470nm signal – fitted signal)/fitted signal)]. Z-scores were calculated using the formula: z-score = ((ΔF/F - mean ΔF/F of baseline period)/standard deviation of ΔF/F of baseline period). For behavior non-specific analysis, the baseline period was 5 to 25 seconds after recording started. For behavior-specific analysis, the baseline period was −10 to −5 seconds before the behavior onset. Area under the curve (AUC) was calculated using the trapezoidal method for specified time points across behaviors. Linear mixed effects models were used to determine the effects of surgery, social condition, and time point using the statsmodels package (94) in Python using one of the following formulas:

- Z-Score=surgery condition + social condition + distance + surgery condition*social condition + surgery condition*distance + social condition*distance + surgery condition*social condition*distance + (1|mouse)
- AUC=surgery condition + time point + surgery condition*time point + (1|mouse)
- AUC=social condition + time point + social condition*time point + (1|mouse)
- AUC=time point + (1|mouse)

## Acknowledgments

Thanks to J. Dong and J. Rashgadol for assistance with fiber photometry. This work was supported by NIH F32MH125597 to ECW, NIH P51OD011107 to ASF, NIH P51OD011107 to AK, NIH U01NS120820 to LT, NSF IOS 1937335, NIH R01MH121829, and a UC Davis Academic Senate Grant to BCT.

## References

1. Goddings A-L, Beltz A, Peper JS, Crone EA, Braams BR (2019): Understanding the Role of Puberty in Structural and Functional Development of the Adolescent Brain. Journal of Research on Adolescence 29: 32–53.

2. Chini M, Hanganu-Opatz IL (2021): Prefrontal Cortex Development in Health and Disease: Lessons from Rodents and Humans. Trends in Neurosciences 44: 227–240.

3. Caballero A, Orozco A, Tseng KY (2021): Developmental regulation of excitatory-inhibitory synaptic balance in the prefrontal cortex during adolescence. Seminars in Cell & Developmental Biology 118: 60–63.

4. Beesdo K, Bittner A, Pine DS, Stein MB, Höfler M, Lieb R, Wittchen H-U (2007): Incidence of social anxiety disorder and the consistent risk for secondary depression in the first three decades of life. Arch Gen Psychiatry 64: 903–912.

5. Dahl RE (2004): Adolescent brain development: a period of vulnerabilities and opportunities. Keynote address. Ann N Y Acad Sci 1021: 1–22.

6. Davey CG, Yücel M, Allen NB (2008): The emergence of depression in adolescence: development of the prefrontal cortex and the representation of reward. Neurosci Biobehav Rev 32: 1–19.

7. Wesselhoeft R, Pedersen CB, Mortensen PB, Mors O, Bilenberg N (2015): Gender–age interaction in incidence rates of childhood emotional disorders. Psychological Medicine 45: 829–839.

8. Hollingworth SA, Burgess PM, Whiteford HA (2010): Affective and anxiety disorders: prevalence, treatment and antidepressant medication use. Aust N Z J Psychiatry 44: 513–519.

9. Hunt C, Issakidis C, Andrews G (2002): DSM-IV generalized anxiety disorder in the Australian National Survey of Mental Health and Well-Being. Psychological Medicine 32: 649–659.

10. Husain N, Creed F, Tomenson B (2000): Depression and social stress in Pakistan. Psychological Medicine 30: 395–402.

11. Ma X, Xiang Y-T, Cai Z-J, Lu J-Y, Li S-R, Xiang Y-Q, et al. (2009): Generalized Anxiety Disorder in China: Prevalence, Sociodemographic Correlates, Comorbidity, and Suicide Attempts. Perspectives in Psychiatric Care 45: 119–127.

12. Munk-Jørgensen P, Allgulander C, Dahl AA, Foldager L, Holm M, Rasmussen I, et al. (2006): Prevalence of Generalized Anxiety Disorder in General Practice in Denmark, Finland, Norway, and Sweden. PS 57: 1738–1744.

13. Remes O, Brayne C, Linde R van der, Lafortune L (2016): A systematic review of reviews on the prevalence of anxiety disorders in adult populations. Brain and Behavior 6: e00497.

14. Dorn LD, Hostinar CE, Susman EJ, Pervanidou P (2019): Conceptualizing Puberty as a Window of Opportunity for Impacting Health and Well-Being Across the Life Span. J Res Adolesc 29: 155–176.

15. Ahmed EI, Zehr JL, Schulz KM, Lorenz BH, DonCarlos LL, Sisk CL (2008): Pubertal hormones modulate the addition of new cells to sexually dimorphic brain regions. Nature Neuroscience 11: 995–997.

16. Delevich K, Piekarski D, Wilbrecht L (2019): Neuroscience: Sex Hormones at Work in the Neocortex. Curr Biol 29: R122–R125.

17. Drzewiecki CM, Willing J, Juraska JM (2016): Synaptic number changes in the medial prefrontal cortex across adolescence in male and female rats: A role for pubertal onset. Synapse 70: 361–368.

18. Schulz KM, Sisk CL (2016): The organizing actions of adolescent gonadal steroid hormones on brain and behavioral development. Neurosci Biobehav Rev 70: 148–158.

19. Delevich K, Thomas AW, Wilbrecht L (2019): Adolescence and “Late Blooming” Synapses of the Prefrontal Cortex. Cold Spring Harb Symp Quant Biol. https://doi.org/10.1101/sqb.2018.83.037507

20. Gegenhuber B, Wu MV, Bronstein R, Tollkuhn J (2022): Gene regulation by gonadal hormone receptors underlies brain sex differences [no. 7912]. Nature 606: 153–159.

21. Guyer AE, Choate VR, Detloff A, Benson B, Nelson EE, Perez-Edgar K, et al. (2012): Striatal functional alteration during incentive anticipation in pediatric anxiety disorders. Am J Psychiatry 169: 205–212.

22. Clauss JA, Blackford JU (2012): Behavioral Inhibition and Risk for Developing Social Anxiety Disorder: A Meta-Analytic Study. Journal of the American Academy of Child & Adolescent Psychiatry 51: 1066-1075.e1.

23. McHenry J, Carrier N, Hull E, Kabbaj M (2014): Sex differences in anxiety and depression: Role of testosterone. Frontiers in Neuroendocrinology 35: 42–57.

24. Bangasser DA, Cuarenta A (2021): Sex differences in anxiety and depression: circuits and mechanisms [no. 11]. Nat Rev Neurosci 22: 674–684.

25. Wellman CL, Bangasser DA, Bollinger JL, Coutellier L, Logrip ML, Moench KM, Urban KR (2018): Sex Differences in Risk and Resilience: Stress Effects on the Neural Substrates of Emotion and Motivation. J Neurosci 38: 9423–9432.

26. Bourke CH, Neigh GN (2011): Behavioral effects of chronic adolescent stress are sustained and sexually dimorphic. Hormones and Behavior 60: 112–120.

27. Asgari P, McKinney G, Hodges TE, McCormick CM (2021): Social Instability Stress in Adolescence and Social Interaction in Female Rats. Neuroscience 477: 1–13.

28. Graf A, Murray SH, Eltahir A, Patel S, Hansson AC, Spanagel R, McCormick CM (2023): Acute and long-term sex-dependent effects of social instability stress on anxiety-like and social behaviours in Wistar rats. Behavioural Brain Research 438: 114180.

29. Hinton EA, Li DC, Allen AG, Gourley SL (2019): Social Isolation in Adolescence Disrupts Cortical Development and Goal-Dependent Decision-Making in Adulthood, Despite Social Reintegration. eNeuro 6: ENEURO.0318-19.2019.

30. Wu X, Arumugam R, Zhang N, Lee MM (2010): Androgen profiles during pubertal Leydig cell development in mice. Reproduction 140: 113–121.

31. Deboer MD, Li Y (2011): Puberty Is Delayed in Male Mice With Dextran Sodium Sulfate Colitis Out of Proportion to Changes in Food Intake, Body Weight, and Serum Levels of Leptin [no. 1]. Pediatr Res 69: 34–39.

32. Sengupta P (2013): The Laboratory Rat: Relating Its Age With Human’s. Int J Prev Med 4: 624–630.

33. Kuske JX, Trainor BC (2021): Mean Girls: Social Stress Models for Female Rodents. Berlin, Heidelberg: Springer, pp 1–30.

34. Steinman MQ, Duque-Wilckens N, Greenberg GD, Hao R, Campi KL, Laredo SA, et al. (2016): Sex-Specific Effects of Stress on Oxytocin Neurons Correspond With Responses to Intranasal Oxytocin. Biological Psychiatry 80: 406–414.

35. Greenberg GD, Laman-Maharg A, Campi KL, Voigt H, Orr VN, Schaal L, Trainor BC (2014): Sex differences in stress-induced social withdrawal: role of brain derived neurotrophic factor in the bed nucleus of the stria terminalis. Frontiers in behavioral neuroscience 7: 223.

36. Duque-Wilckens N, Torres LY, Yokoyama S, Minie VA, Tran AM, Petkova SP, et al. (2020): Extra-hypothalamic oxytocin neurons drive stress-induced social vigilance and avoidance. Proc Natl Acad Sci USA 117: 26406–26413.

37. Duque-Wilckens N, Steinman MQ, Busnelli M, Chini B, Yokoyama S, Pham M, et al. (2018): Oxytocin Receptors in the Anteromedial Bed Nucleus of the Stria Terminalis Promote Stress-Induced Social Avoidance in Female California Mice. Biological psychiatry 83: 203–213.

38. Luo PX, Zakharenkov HC, Torres LY, Rios RA, Gegenhuber B, Black AM, et al. (2022): Oxytocin receptor behavioral effects and cell types in the bed nucleus of the stria terminalis. Hormones and Behavior 143: 105203.

39. Trainor BC, Pride MC, Landeros RV, Knoblauch NW, Takahashi EY, Silva AL, Crean KK (2011): Sex differences in social interaction behavior following social defeat stress in the monogamous California mouse (peromyscus californicus). PLoS ONE 6. https://doi.org/10.1371/journal.pone.0017405

40. Trainor BC, Takahashi EY, Campi KL, Florez SA, Greenberg GD, Laman-Maharg A, et al. (2013): Sex differences in stress-induced social withdrawal: independence from adult gonadal hormones and inhibition of female phenotype by corncob bedding. Hormones and Behavior 63: 543–550.

41. Greenberg GD, Steinman MQ, Doig IE, Hao R, Trainor BC (2015): Effects of social defeat on dopamine neurons in the ventral tegmental area in male and female California mice. The European journal of neuroscience 42: 3081–94.

42. Duque-Wilckens N, Steinman MQ, Busnelli M, Chini B, Yokoyama S, Pham M, et al. (2018): Oxytocin Receptors in the Anteromedial Bed Nucleus of the Stria Terminalis Promote Stress-Induced Social Avoidance in Female California Mice. Biol Psychiatry 83: 203–213.

43. Numa C, Nagai H, Taniguchi M, Nagai M, Shinohara R, Furuyashiki T (2019): Social defeat stress-specific increase in c-Fos expression in the extended amygdala in mice: Involvement of dopamine D1 receptor in the medial prefrontal cortex. Sci Rep 9: 16670.

44. Kollack-Walker, Don, Watson, Akil (1999): Differential Expression of c-*fos* mRNA Within Neurocircuits of Male Hamsters Exposed to Acute or Chronic Defeat. Journal of Neuroendocrinology 11: 547–559.

45. Samonds JM, Choi V, Priebe NJ (2019): Mice Discriminate Stereoscopic Surfaces Without Fixating in Depth. J Neurosci 39: 8024–8037.

46. Lewis EM, Stein-O’Brien GL, Patino AV, Nardou R, Grossman CD, Brown M, et al. (2020): Parallel Social Information Processing Circuits Are Differentially Impacted in Autism. Neuron 108: 659-675.e6.

47. Trainor BC, Marler CA (2002): Testosterone promotes paternal behaviour in a monogamous mammal via conversion to oestrogen. Proceedings of the Royal Society of London Series B: Biological Sciences 269: 823–829.

48. Hostinar CE, Johnson AE, Gunnar MR (2015): Parent support is less effective in buffering cortisol stress reactivity for adolescents compared to children. Dev Sci 18: 281–297.

49. Wright EC, Hostinar CE, Trainor BC (2020): Anxious to see you: Neuroendocrine mechanisms of social vigilance and anxiety during adolescence. European Journal of Neuroscience 52: 2516–2529.

50. Trainor BC, Pride MC, Villalon Landeros R, Knoblauch NW, Takahashi EY, Silva AL, Crean KK (2011): Sex Differences in Social Interaction Behavior Following Social Defeat Stress in the Monogamous California Mouse (<italic>Peromyscus californicus</italic>). PLoS ONE 6: e17405.

51. Kollack-Walker S, Newman SW (1995): Mating and agonistic behavior produce different patters of Fos immunolabeling in the male Syrian hamster brain. Neuroscience 66: 721– 736.

52. Nasanbuyan N, Yoshida M, Takayanagi Y, Inutsuka A, Nishimori K, Yamanaka A, Onaka T (2018): Oxytocin-Oxytocin Receptor Systems Facilitate Social Defeat Posture in Male Mice. Endocrinology 159: 763–775.

53. Newman EL, Covington HE, Suh J, Bicakci MB, Ressler KJ, DeBold JF, Miczek KA (2019): Fighting Females: Neural and Behavioral Consequences of Social Defeat Stress in Female Mice. Biol Psychiatry 86: 657–668.

54. Martinez M, Phillips PJ, Herbert J (1998): Adaptation in patterns of *c-fos* expression in the brain associated with exposure to either single or repeated social stress in male rats. European Journal of Neuroscience 10: 20–33.

55. Fox AS, Oler JA, Shackman AJ, Shelton SE, Raveendran M, McKay DR, et al. (2015): Intergenerational neural mediators of early-life anxious temperament. Proc Natl Acad Sci USA 112: 9118–9122.

56. Somerville LH, Wagner DD, Wig GS, Moran JM, Whalen PJ, Kelley WM (2013): Interactions Between Transient and Sustained Neural Signals Support the Generation and Regulation of Anxious Emotion. Cerebral Cortex 23: 49–60.

57. Davis M, Walker DL, Miles L, Grillon C (2010): Phasic vs Sustained Fear in Rats and Humans: Role of the Extended Amygdala in Fear vs Anxiety. Neuropsychopharmacology 35: 105– 135.

58. Shackman AJ, Fox AS (2016): Contributions of the central extended amygdala to fear and anxiety. J Neurosci 36: 8050–8063.

59. Steinman MQ, Laredo SA, Lopez EM, Manning CE, Hao RC, Doig IE, et al. (2015): Hypothalamic vasopressin systems are more sensitive to the long term effects of social defeat in males versus females. Psychoneuroendocrinology 51. https://doi.org/10.1016/j.psyneuen.2014.09.009

60. Flanigan ME, Kash TL (2020): Coordination of social behaviors by the bed nucleus of the stria terminalis. European Journal of Neuroscience in press. https://doi.org/10.1111/ejn.14991

61. Lebow MA, Chen A (2016): Overshadowed by the amygdala: the bed nucleus of the stria terminalis emerges as key to psychiatric disorders. Mol Psychiatry 21: 450–463.

62. Lissek S, Powers AS, McClure EB, Phelps EA, Woldehawariat G, Grillon C, Pine DS (2005): Classical fear conditioning in the anxiety disorders: a meta-analysis. Behaviour Research and Therapy 43: 1391–1424.

63. Fox AS, Shelton SE, Oakes TR, Davidson RJ, Kalin NH (2008): Trait-Like Brain Activity during Adolescence Predicts Anxious Temperament in Primates. PLOS ONE 3: e2570.

64. Clauss JA, Avery SN, Benningfield MM, Blackford JU (2019): Social anxiety is associated with BNST response to unpredictability. Depress Anxiety 36: 666–675.

65. Williams ES, Manning CE, Eagle AL, Swift-Gallant A, Duque-Wilckens N, Chinnusamy S, et al. (2020): Androgen-Dependent Excitability of Mouse Ventral Hippocampal Afferents to Nucleus Accumbens Underlies Sex-Specific Susceptibility to Stress. Biol Psychiatry 87: 492–501.

66. Harley CW, Malsbury CW, Squires A, Brown RA (2000): Testosterone decreases CA1 plasticity in vivo in gonadectomized male rats. Hippocampus 10: 693–697.

67. Hebbard PC, King RR, Malsbury CW, Harley CW (2003): Two organizational effects of pubertal testosterone in male rats: transient social memory and a shift away from long-term potentiation following a tetanus in hippocampal CA1. Experimental Neurology 182: 470–475.

68. Moradpour F, Fathollahi Y, Naghdi N, Hosseinmardi N, Javan M (2013): Prepubertal castration causes the age-dependent changes in hippocampal long-term potentiation. Synapse 67: 235–244.

69. Yu W, Caira CM, Del R Rivera Sanchez N, Moseley GA, Kash TL (2021): Corticotropin-releasing factor neurons in the bed nucleus of the stria terminalis exhibit sex-specific pain encoding in mice. Sci Rep 11: 12500.

70. Rodriguez-Romaguera J, Ung RL, Nomura H, Otis JM, Basiri ML, Namboodiri VMK, et al. (2020): Prepronociceptin-Expressing Neurons in the Extended Amygdala Encode and Promote Rapid Arousal Responses to Motivationally Salient Stimuli. Cell Reports 33: 108362.

71. Yang B, Karigo T, Anderson DJ (2022): Transformations of neural representations in a social behaviour network [no. 7924]. Nature 608: 741–749.

72. Kaouane N, Ada S, Hausleitner M, Haubensak W (2021): Dorsal Bed Nucleus of the Stria Terminalis-Subcortical Output Circuits Encode Positive Bias in Pavlovian Fear and Reward. Front Neural Circuits 15: 772512.

73. Tian G, Hui M, Macchia D, Derdeyn P, Rogers A, Hubbard E, et al. (2022): An extended amygdala-midbrain circuit controlling cocaine withdrawal-induced anxiety and reinstatement. Cell Reports 39: 110775.

74. Kim S-Y, Adhikari A, Lee SY, Marshel JH, Kim CK, Mallory CS, et al. (2013): Diverging neural pathways assemble a behavioural state from separable features in anxiety [no. 7444]. Nature 496: 219–223.

75. Jennings JH, Sparta DR, Stamatakis AM, Ung RL, Pleil KE, Kash TL, Stuber GD (2013): Distinct extended amygdala circuits for divergent motivational states. Nature 496: 224–228.

76. Giardino WJ, Eban-Rothschild A, Christoffel DJ, Li S-B, Malenka RC, de Lecea L (2018): Parallel circuits from the bed nuclei of stria terminalis to the lateral hypothalamus drive opposing emotional states [no. 8]. Nat Neurosci 21: 1084–1095.

77. Cooper MA, Clinard CT, Dulka BN, Grizzell JA, Loewen AL, Campbell AV, Adler SG (2021): Gonadal steroid hormone receptors in the medial amygdala contribute to experience-dependent changes in stress vulnerability. Psychoneuroendocrinology 129: 105249.

78. Aikey JL, Nyby JG, Anmuth DM, James PJ (2002): Testosterone Rapidly Reduces Anxiety in Male House Mice (Mus musculus). Hormones and Behavior 42: 448–460.

79. Frye CA, Seliga AM (2001): Testosterone increases analgesia, anxiolysis, and cognitive performance of male rats. Cognitive, Affective, & Behavioral Neuroscience 1: 371–381.

80. Carrier N, Kabbaj M (2012): Extracellular Signal-Regulated Kinase 2 Signaling in the Hippocampal Dentate Gyrus Mediates the Antidepressant Effects of Testosterone. Biological Psychiatry 71: 642–651.

81. Seeman TE, Singer B, Wilkinson CW, Bruce McEwen (2001): Gender differences in age-related changes in HPA axis reactivity. Psychoneuroendocrinology 26: 225–240.

82. Uhart M, Chong R, Oswald L, Lin P, Wand G (2006): Gender differences in hypothalamic– pituitary–adrenal (HPA) axis reactivity. Psychoneuroendocrinology 31: 642–652.

83. Kirschbaum C, Kudielka BM, Gaab J, Schommer NC, Hellhammer DH (1999): Impact of Gender, Menstrual Cycle Phase, and Oral Contraceptives on the Activity of the Hypothalamus-Pituitary-Adrenal Axis: Psychosomatic Medicine 61: 154–162.

84. Borrow AP, Handa RJ (2017): Estrogen Receptors Modulation of Anxiety-Like Behavior. Vitamins and Hormones, vol. 103. Elsevier, pp 27–52.

85. Kraynak M, Flowers MT, Shapiro RA, Kapoor A, Levine JE, Abbott DH (2017): Extraovarian gonadotropin negative feedback revealed by aromatase inhibition in female marmoset monkeys. Am J Physiol Endocrinol Metab 313: E507–E514.

86. Collins HH (1923): Studies of the pelage phases and of the nature of color variations in mice of the genus Peromyscus. Journal of Experimental Zoology 38: 45–107.

87. Ounpraseuth S, Rafferty TM, McDonald-Phillips RE, Gammill WM, Siegel ER, Wheeler KL, et al. (2009): A Method to Quantify Mouse Coat-Color Proportions. PLOS ONE 4: e5414.

88. Murphy KL, Baxter MG (2013): Long-Term Effects of Neonatal Single or Multiple Isoflurane Exposures on Spatial Memory in Rats. Front Neurol 4. https://doi.org/10.3389/fneur.2013.00087

89. Lauer J, Zhou M, Ye S, Menegas W, Nath T, Rahman MM, et al. (2021): Multi-Animal Pose Estimation and Tracking with DeepLabCut. Animal Behavior and Cognition. https://doi.org/10.1101/2021.04.30.442096

90. Nath T, Mathis A, Chen AC, Patel A, Bethge M, Mathis MW (2019): Using DeepLabCut for 3D markerless pose estimation across species and behaviors. Nat Protoc 14: 2152–2176.

91. Mathis A, Mamidanna P, Cury KM, Abe T, Murthy VN, Mathis MW, Bethge M (2018): DeepLabCut: markerless pose estimation of user-defined body parts with deep learning. Nat Neurosci 21: 1281–1289.

92. Trainor BC, Bird IM, Alday NA, Schlinger BA, Marler CA (2003): Variation in Aromatase Activity in the Medial Preoptic Area and Plasma Progesterone Is Associated with the Onset of Paternal Behavior. NEN 78: 36–44.

93. Minie VA, Petric R, Ramos-Maciel S, Wright EC, Trainor BC, Duque-Wilckens N (2021): Enriched laboratory housing increases sensitivity to social stress in female California mice (Peromyscus californicus). Applied Animal Behaviour Science 241: 105381.

94. Seabold S, Perktold J (2010): Statsmodels: Econometric and Statistical Modeling with Python. 92–96.

