## Supplemental Figures for "Sexual differentiation of neural mechanisms of stress sensitivity during puberty"

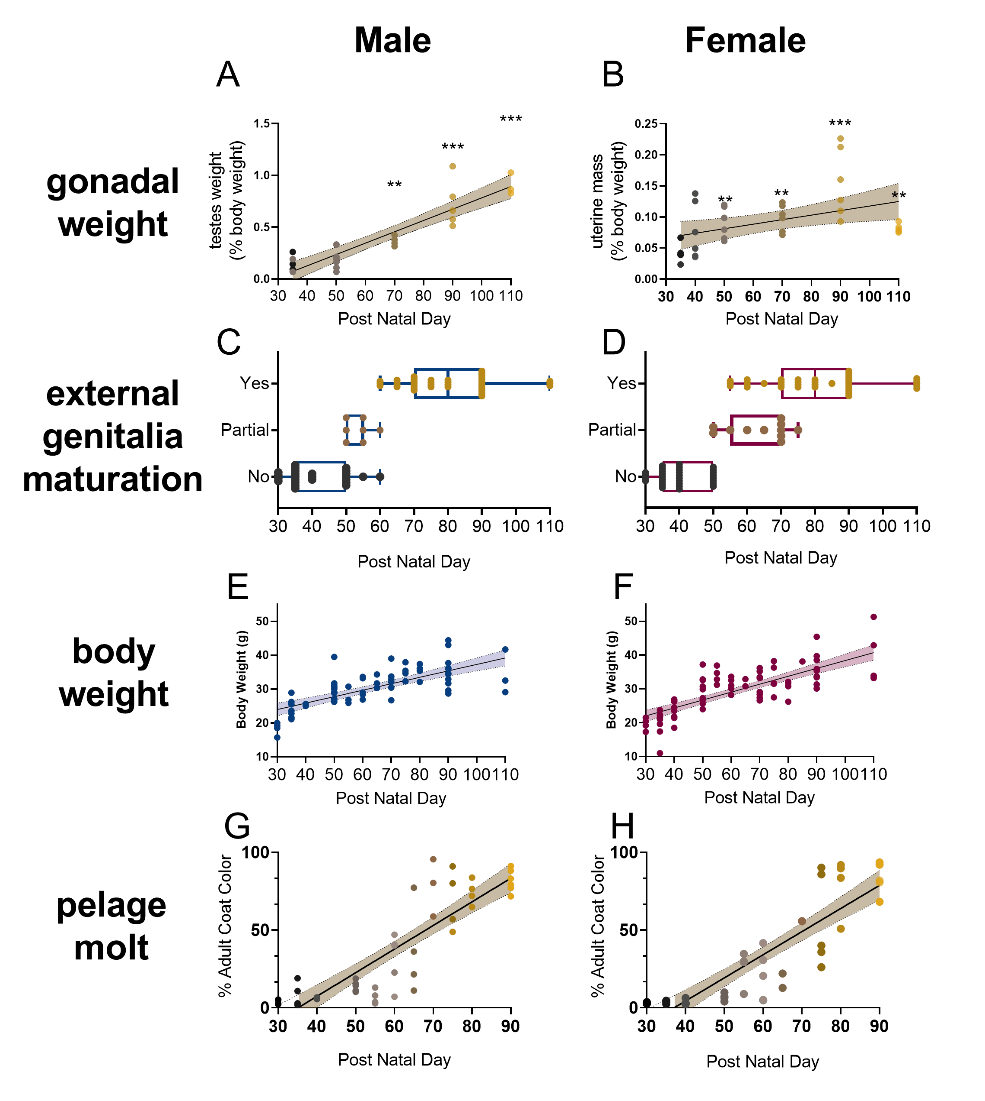

**Supplementary figure 1**: Metrics of pubertal development in male and female California mice. Male testes weight (A) and female uterine (B) weight increases with age. Mice of different ages were assessed for preputial separation (C) or vaginal opening (D) to reveal that external genitalia maturation generally occurs after post-natal day 60. Measures of body weight in male (E) and female (F) mice increases with age. California mice molt from a gray pelage to a brown pelage, with the fastest rate of change occurring after post-natal day 60. ** p<0.01, *** p<0.001 versus post-natal day 35.

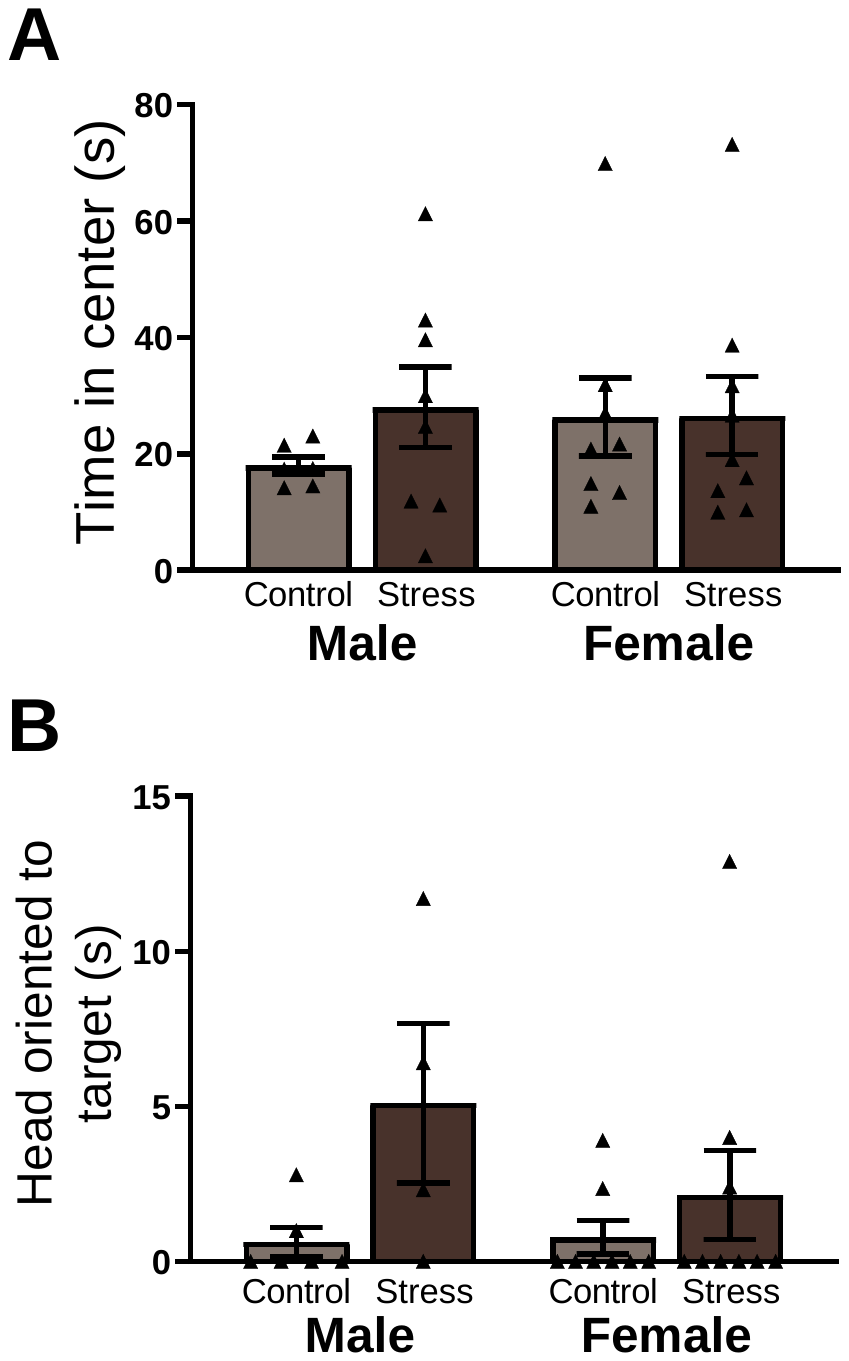

**Supplementary figure 2**: For juvenile mice there were no differences in time spent in the center of the arena during open field (A) or in vigilance during the acclimation phase (B).

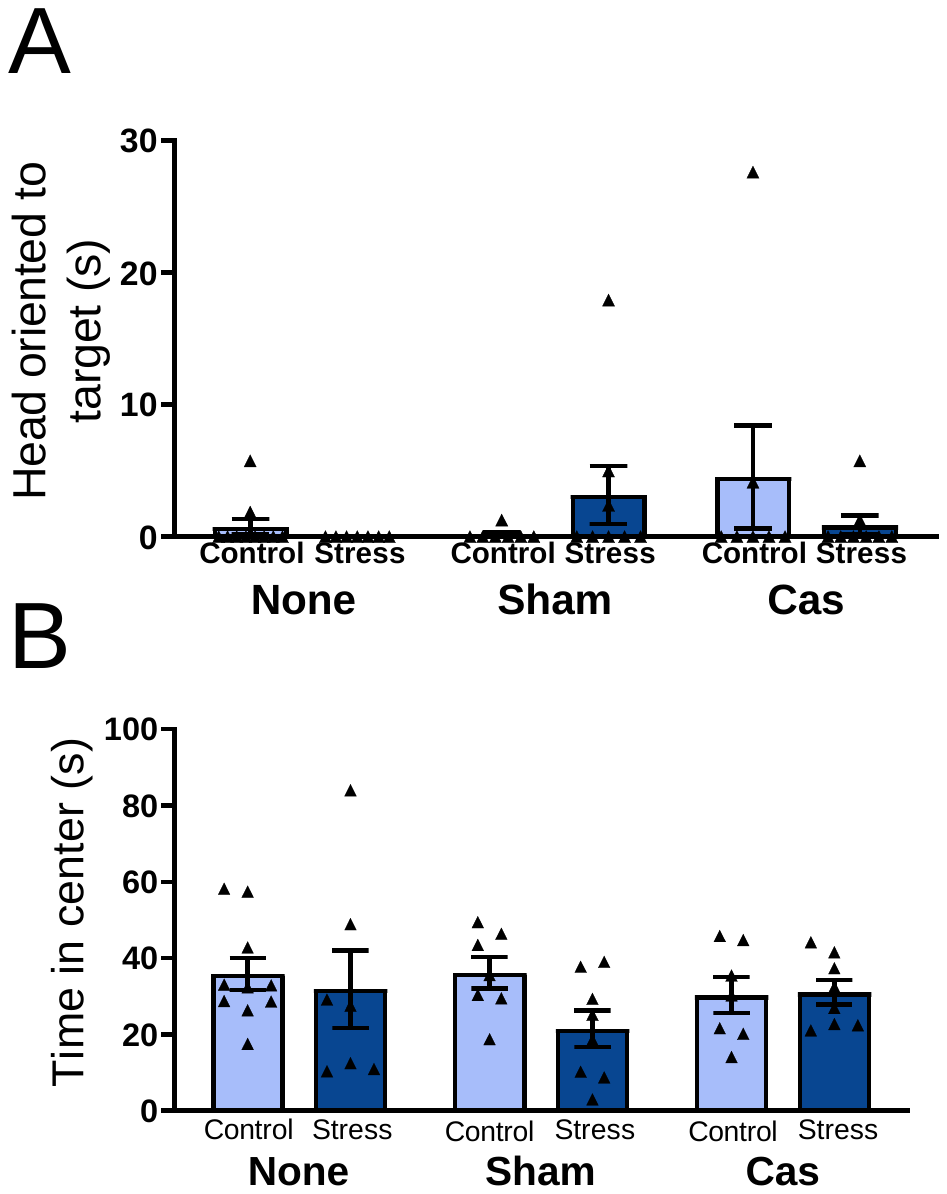

**Supplementary figure 3**: For adult male mice, there were no differences in vigilance during the acclimation phase (A) or in time spent in the center of the arena during the open field phase (B).

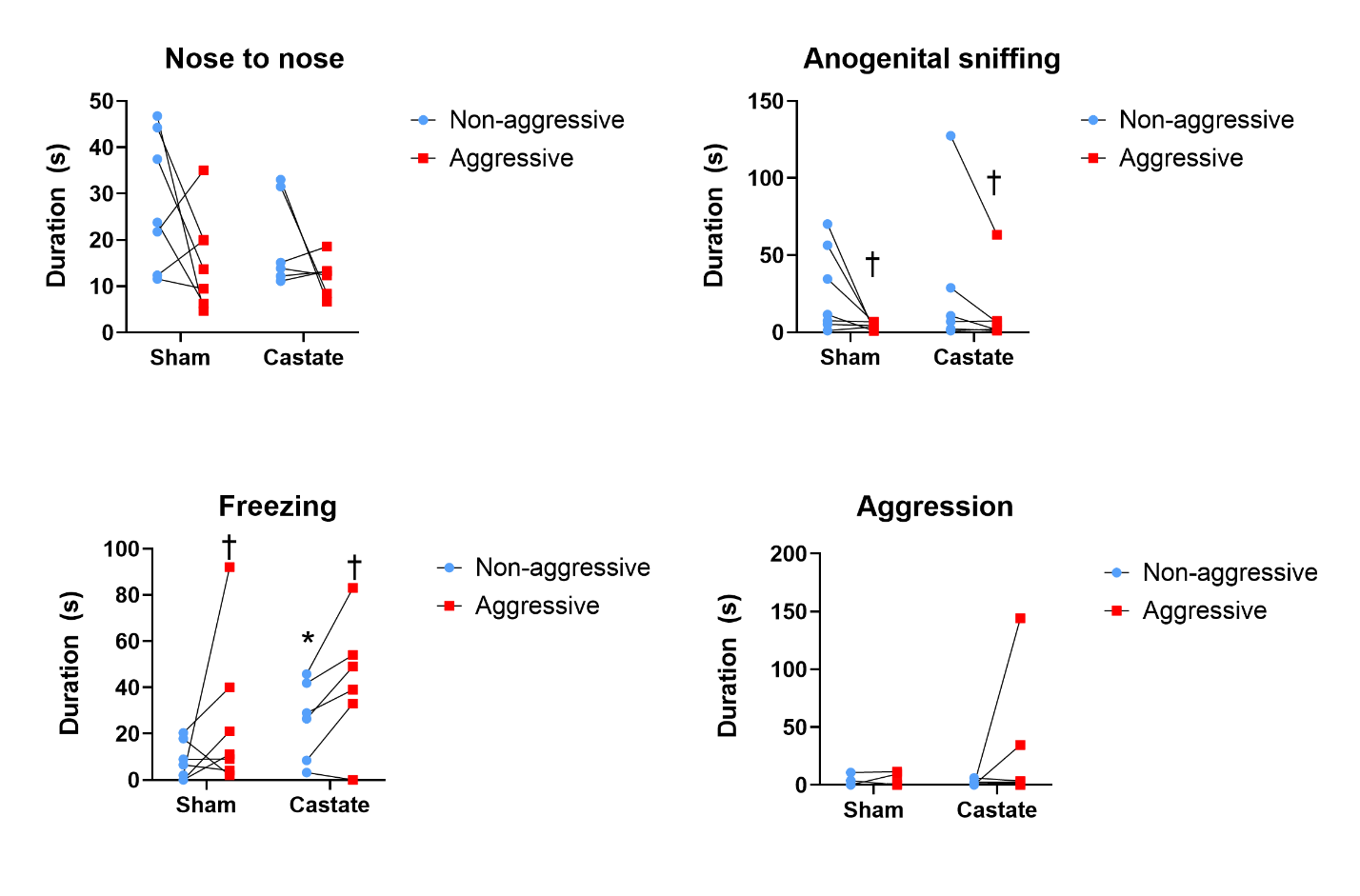

**Supplementary figure 4**: Behavior of sham and prepubertal castrated male mice during photometry observations with non-aggressive and aggressive target mice. * p <0.05 vs. sham. † p<0.05 vs non-aggressive target mouse.

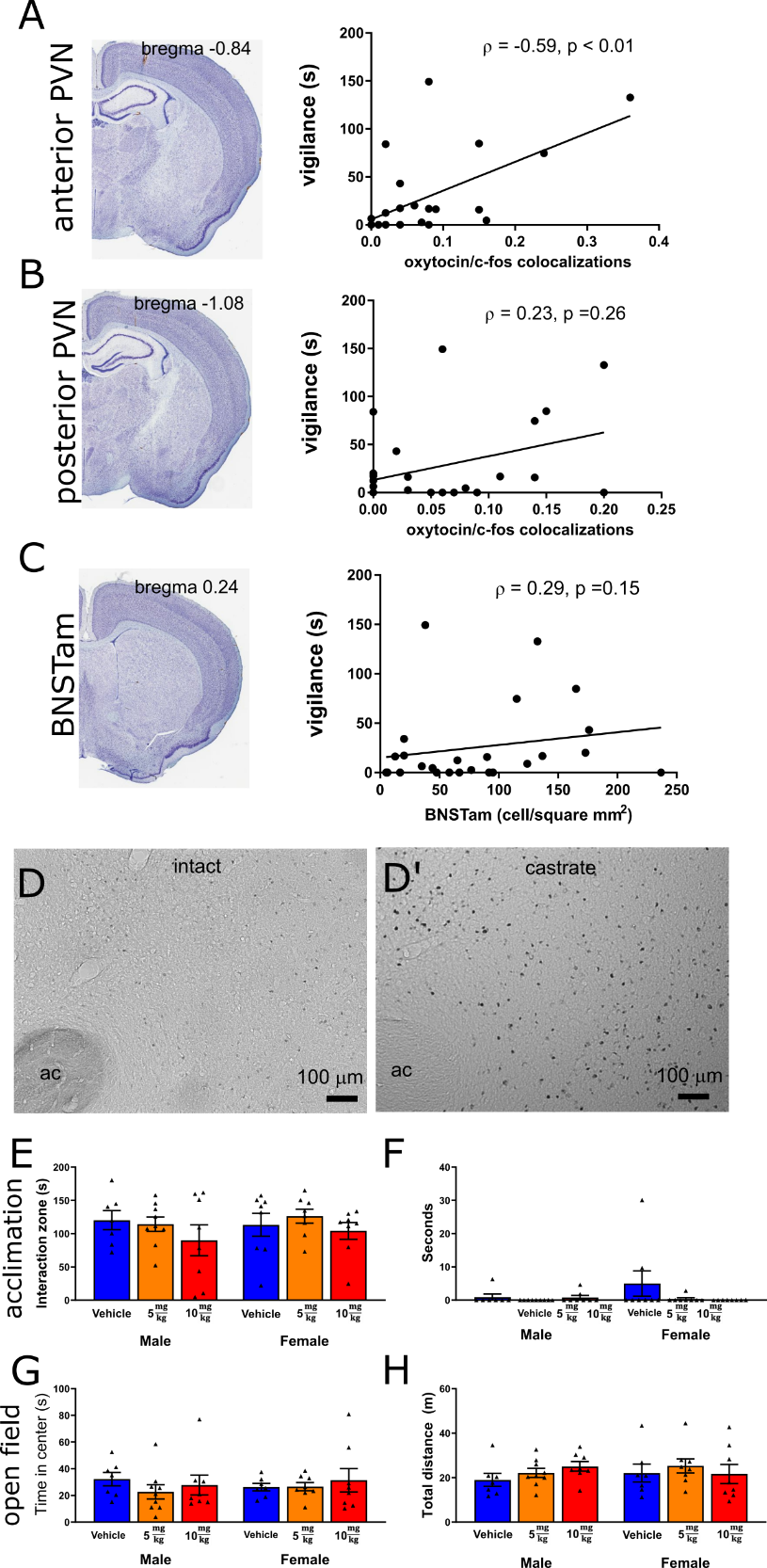

**Supplementary figure 5**: Social vigilance was positively correlated with oxytocin/c-fos colocalizations in the anterior (A) but not posterior (B) PVN, and did not correlate with c-fos in anteromedial BNST (C). Castration increased c-fos immunoreactivity in the anteromedial BNST (D, D’). Oxytocin receptor antagonists did not affect time in the interaction zone (E) or vigilance (F) during the acclimation stage and did not affect time in the center (G) or locomotor behavior (H) during the open field phase.

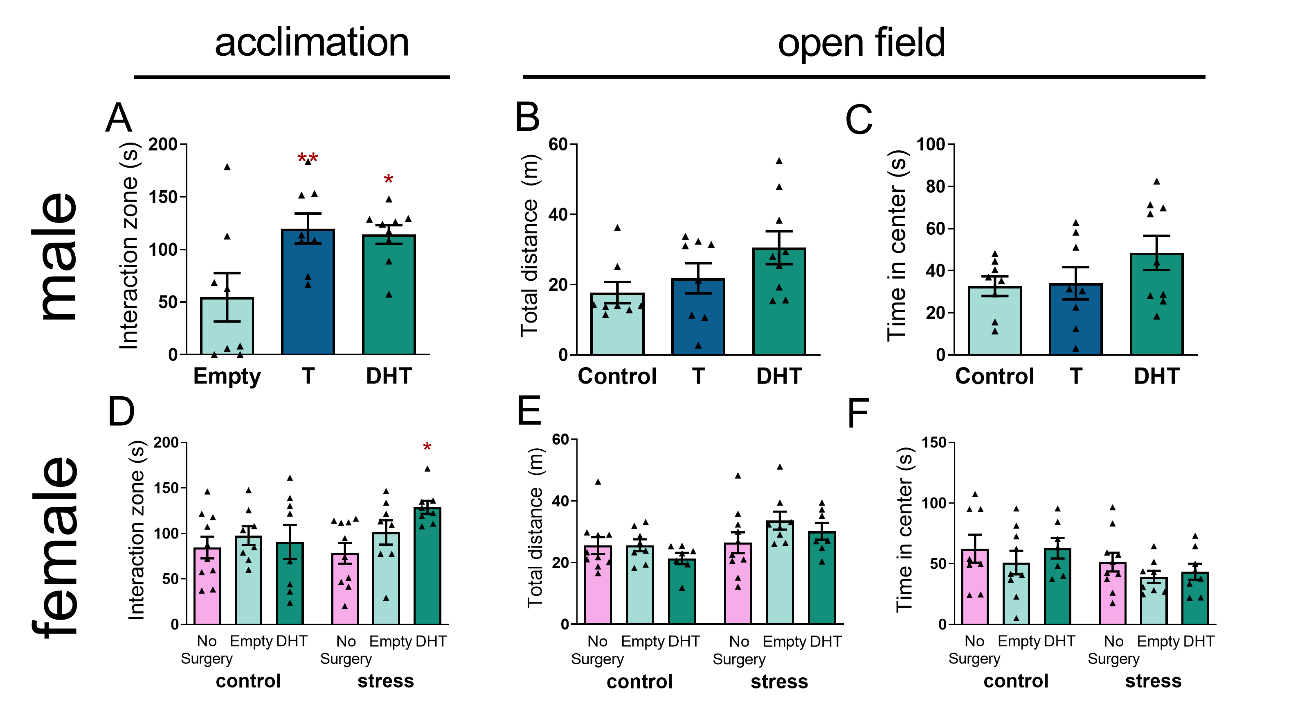

**Supplementary figure 6**: In stressed males, T and DHT treatment increased approach to an empty cage (A) but had not effects on locomotor behavior (B) or time in the center (C) during the open field phase. For females, DHT treatment had no effect on approach to an empty cage (D) or behavior in the open field phase (E,F). After stress DHT increased approach to an empty cage. * p<0.05, **p<0.01 vs empty implant.
